# Fin sweep angle does not determine flapping propulsive performance

**DOI:** 10.1101/2020.11.03.366997

**Authors:** Andhini N. Zurman-Nasution, Bharathram Ganapathisubramani, Gabriel D. Weymouth

## Abstract

The importance of the leading-edge sweep angle of propulsive surfaces used by unsteady swimming and flying animals has been an issue of debate for many years, spurring studies in biology, engineering, and robotics with mixed conclusions. In this work, we provide results from three-dimensional simulations of finite foils undergoing tail-like (pitch-heave) and flipper-like (twist-roll) kinematics for a range of sweep angles based on a large range of animal data while carefully controlling all other parameters. Our primary finding is the negligible 0.043 maximum correlation between the sweep angle and the propulsive force and power for both tail-like and flipper-like motions. This indicates that fish tails and mammal flukes with similar range and size can have a large range of potential sweep angles without significant negative propulsive impact. Although there is a slight benefit to *avoiding* large sweep angles, this is easily compensated by adjusting the fin’s motion parameters such as flapping frequency, amplitude and maximum angle of attack to gain higher thrust and efficiency.

## Introduction

A large fraction of swimming and flying animals use propulsive flapping to move, and there are significant potential applications for this biologically inspired form of propulsion in engineering and robotics. As such, significant research effort has been devoted towards understanding and optimizing both the kinematics and morphology of the propulsion surface itself, with a particular emphasis on explaining propulsive efficiency.

The leading edge sweep back angle is a key geometric feature observed in fish tails, mammal flukes, aquatic-animal flippers, and bird and insect wings. There have been extensive studies collecting sweep back angle data from a variety of animal propulsion surfaces. The data in Fig. 1 are collected from marine mammals (1), fish caudal fins (2), flyer wings (3–7), and aquatic-animal propulsive side-fins such as flippers (8–11), pectoral (12, 13) and dorsal fins (14). There is a strong correlation between fin aspect ratio, 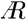, and kinematics. Most tails in red colors (mammal flukes and caudal fins) are limited to the low to medium 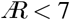 fins, where their median and average value are around 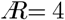. The highest 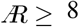 region is populated only by flippers and wings which performing rolling and twisting motions, such as penguins, turtle and most plesiosaurs. In contrast, the sweep angle Λ dependence is much less clear, with flipper and propulsive pectoral fins spanning a wide range of angles. While some individual studies indicate that increased sweep angles are correlated with animals with lower aspect ratios 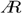 fins (15, 16), there is extensive variation between animals within the groups and no strong trend is found when including all the groups using propulsive flapping. Additionally, there are hundreds of differences in the specific geometry and kinematics of each of these animals and just as many biological pressures other than propulsive efficiency at play, making it difficult to conclusively determine the influence of sweep angle from biological data directly.

**Fig. 1.**
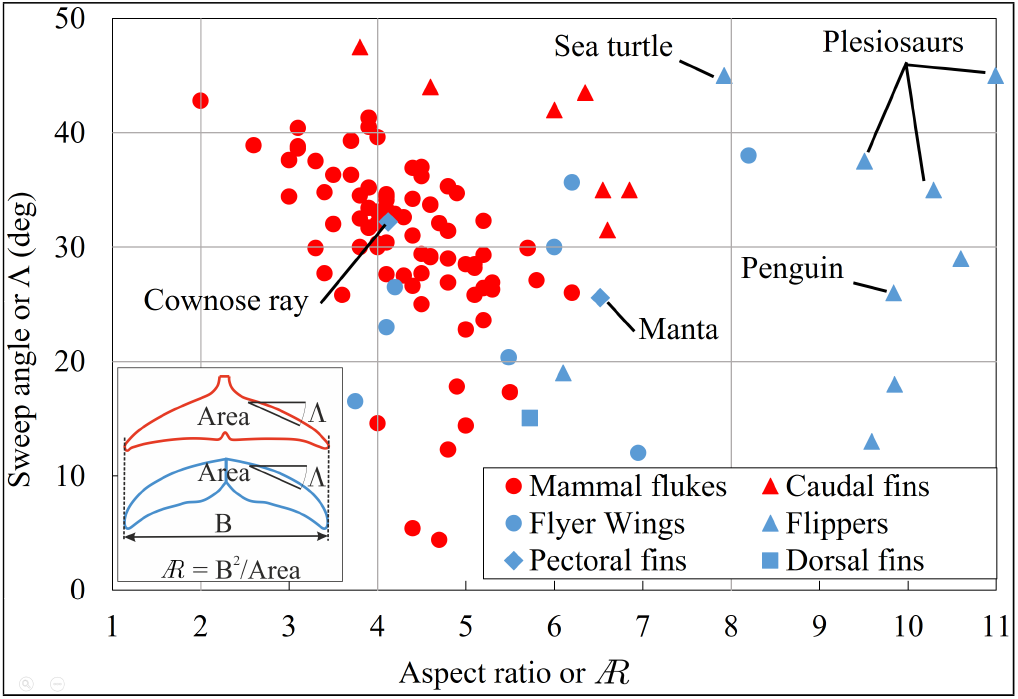
Collection of biologist data for sweep angles Λ and aspect ratio 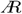 for fish caudal fins and mammal flukes (in red) and wings, flippers, pectoral, and dorsal fins wings (in blue) shows the extremely wide scatter across this evolutionary parameter space for different successful animals. The current computational study takes place over sweep angles from 20, 30&40 deg and 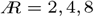 shown by grey lines to maximize the relevance to biological propulsive surfaces.

Focused engineering studies have also been applied to determine the impact of sweep angle on the performance of flapping propulsion surfaces. While these would hopefully clarify the issues at play, the finding are somewhat contradictory, especially between theoretical and experimental analysis. Most analytical studies using lifting-line theory indicate sweep is advantageous, increasing the thrust and propulsive efficiency (15–17). While such methodologies are effective at determining lift characteristics on steady wings, they cannot model unsteady rotational three-dimensional flow, including the evolution of vortices that form at the fin’s leading edge and tip which lead to the high forces observed in flapping foil propulsion (18, 19).

While some experimental studies have measured a small flapping propulsive benefit to sweep, the effect is smaller than the analytic studies and other experiments have found no impact at all. A study on the spanwise flow caused by sweep angle found that it did not stabilizing the LEV on an impulsively heaved foil (20), in contrast to the known stabilizing effect on a stationary wing. Another study reported affected forces and vortex strength on a 45°-sweptback flapping plate for large-scale LEVs produced by an impulsively heaved plate (21). Insignificant force change is also seen in the experiment of manta robot with rolling flexible flippers (22). Varying sweep angle from zero to 60° in rotating wings found no influence on the stability of the LEV (23). For a purely pitching flat plate, the data of circulation time-histories show varying reduced frequencies can change the LEV circulation slightly more on the lower frequency cases (24). This indicates that it is important to test a variety of kinematics relevant to the problem, as well as controlling for as many other parameters as possible, something which is very difficult to accomplish in physical experiments.

In this work, we carefully isolate the effect of varied sweep angles from all other geometric parameters to help address the conflicting and scattered results of previous biological, theoretical and experimental studies. Using 3D unsteady numerical simulations, we add sweep to a simple base fin shape and test its flapping propulsive performance for two types of biologically inspired kinematics, i.e. tail-like and flipper-like motions, Fig. 2(a). We also test the combined impact of 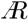 and sweep as well as other kinematic parameters such as motion types and amplitudes to determine the relative influence of sweep angle on propulsive flapping.

**Fig. 2.**
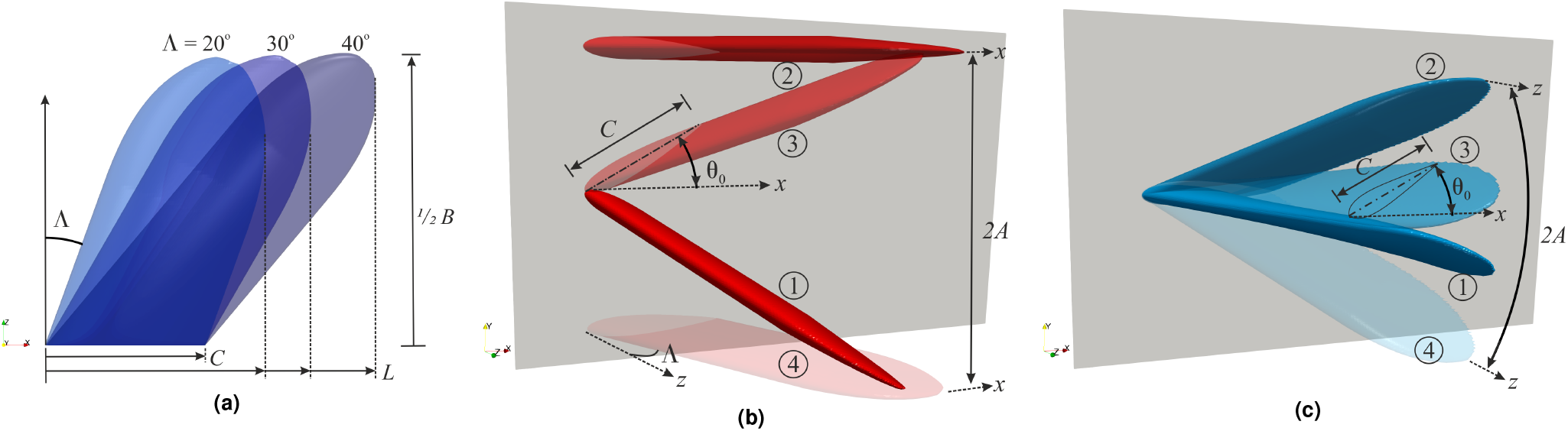
Diagram of the propulsive fin geometry and kinematics. (a) Foil planform with various sweep angle Λ for two different kinematic i.e. (b) fishtail-like kinematics i.e. pitch-heave combined with 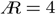, and (c) flipper-like kinematics i.e. twist-roll combined with 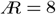. Phase steps per cycle are denoted from ➀ to ➃.

Specifically, we define our geometry to use a NACA0016 foil cross-section with chord length *C*, defined parallel to the inflow the *U*_∞_, and thickness *D* = 0.16*C*. A rectangular planform is combined with a tapered elliptic tip with a length of 1*C*. The sweep angle Λ, defined in Fig. 2, is varied from Λ = [20°, 30°, 40°] to cover the most populated biological range, Fig. 1. As the sweep is adjusted, the tip-to-tip foil span *B* is adjusted to maintain the desired aspect ratio 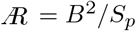, where *S*_*p*_ is the planform area. As with sweep, we use a wide range of 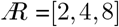 to cover the 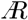 range of propulsive surfaces found in nature. Our chosen ranges have covered a minimum of 95% for aspect-ratio and 82% for sweep angle as well as the characteristics *Re* range from the population of animal data provided in Fig.1. The chord-wise Reynolds number *Re* = *ρU*_∞_*C/µ* = 5300, where *ρ, µ* are the fluid density and viscosity, is kept fixed to isolate influence of the geometry and kinematics. This Reynolds number is representative for a wide range of biological and robotic applications as the thrust and power coefficients of flapping foils at the same Strouhal number and similar offset drag are insensitive to the variation of chordwise Reynolds number between *Re ≈*5000 and *Re ≈*500, 000 (25, 26).

This geometry is subjected to sinusoidal “tail-like” and “flipper-like” motions diagrammed in Fig. 2. The tail-like kinematics are inspired by biologist data for fish caudal fins and mammal flukes and made up coupled heave and pitch motions, Fig 2(b). The flipper-like kinematics are inspired by aquatic animal pectoral flippers and bird/insect wings, and is made up twist and roll motions, Fig 2(c). Combining twist to roll acts as prescribed linear flexibility to rigid foil. The motion amplitude and frequency are quantified by Strouhal number *St* = 2*Af/U*_∞_ and reduced frequency *k*^*^ = *fl*^*^*/U*_∞_, where *f* is the motion frequency, *A* is the total perpendicular amplitude envelop for the motion, and *l*^*^ is a length-scale equal to the total inline extent *L* for tail-like motions and *B/*2 for flipper-like motions. The majority of the simulations use a fairly low frequency *k*^*^ = 0.3 and high amplitude 2*A* = *l*^*^ condition to imitate the average mammal-fluke kinematics as reported in biological data (1), and the Strouhal number *St* = 0.3 is fixed in the middle of the optimal propulsive efficiency range (27). A few special cases outside these conditions were also simulated: A flipper-like motion using higher *k*^*^ = 0.6 and *St* = 0.6 to imitate penguin and turtle swimming data with high roll and twist angles to maintain high propulsion and efficiency due to high 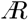. Also, a tail-like motion at lower amplitude *A* = 0.13*l*^*^ and much higher frequency at *St* = 0.46 inspired by recent robotics studies at similar operating conditions (28, 29).

### Impact of sweep on wake structures and propulsive characteristics

A set of example simulation results for both tail-like and flipper-like motions are shown in Fig. 3, with the flow’s vortex structures visualized using the *Q*-criterion (29). The complete unsteady flow field evolution can be viewed in the supplementary video. These wake visualizations show that small scale features such as the generation of small turbulent vortices depends on both the geometry and the kinematics. However, the large-scale underlying flow structures illustrated in the bottom row sketches are the same for a given set of kinematics; changing the sweep angle from 20° to 40° simply shifts and scales those structures. Specifically, for the tail-like motions in the left column, the leading edge vortex detaches from the top of the fin halfway along the inline length *l*^*^ = *L* for tail-like motions. Similarly, while increasing the sweep angle pushes the LEV back for flipper like motions, the separation for both cases occurs around 70% along the fin width *l*^*^ = *B/*2 for these cases. The period and wavelength of the trailing wake structures also scale with *l*^*^ because we have used *l*^*^ to define the reduced frequency *k\** and the amplitude envelop 2*A* = *l*^*^. In other words, the sweep angle does not fundamentally change the resulting propulsive flow around the fins other than changing the characteristic length *l*^*^.

**Fig. 3.**
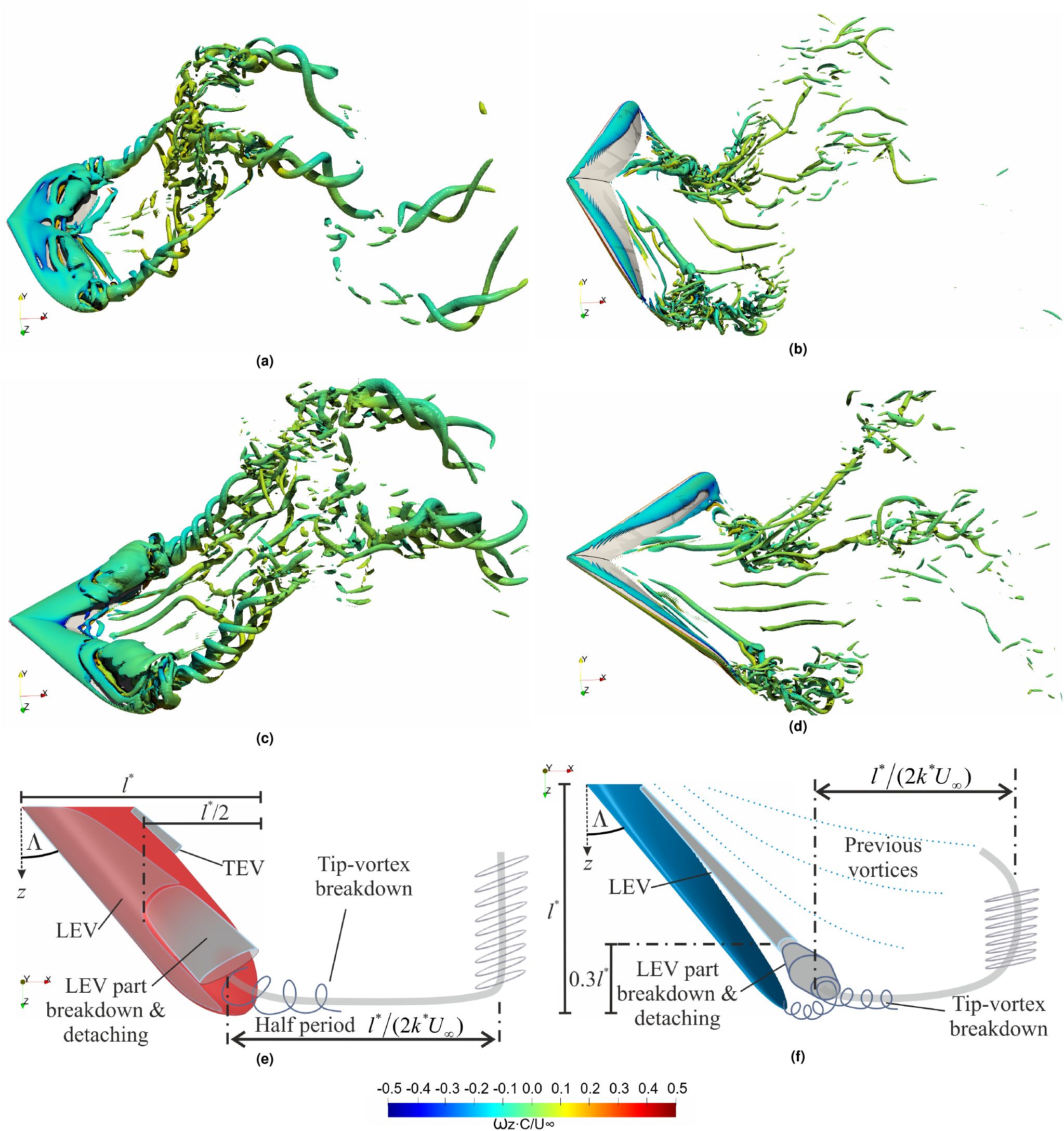
Flow structures generated by tail-like and flipper-like propulsive swimming at position ➀ shown in Fig.2. Tail-like motions are depicted in pitch-heave for (a) Λ20, (c) Λ40 for 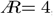, and flipper-like motions as in twist-roll for (b) Λ20, (d) Λ40 for 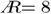. Instantaneous vortex structures are visualized using iso-surfaces with 1% of *Q*_*max*_ colored by the spanwise vorticity *ω*_*z*_ *C/U*_*∞*_. Illustration of vortex-shedding process every half period (*l*^*\**^ */*(2*k*^*\**^ *U*_*∞*_)) of cycle for (e) tail-like motion and (f) flipper-like.

We can quantify this invariance of the flow in terms of the integrated propulsive metrics; the coefficients of thrust *C*_*T*_ lift *C*_*L*_ and power *C*_*Pow*_, and the propulsive efficiency *η*, defined as

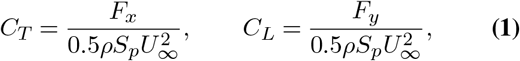

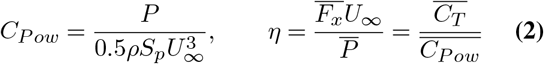

where measured thrust force (*F*_*x*_), lift force (*F*_*y*_), and power (*P*) are calculated from the integration of pressure and viscous forces over the foil. *S*_*p*_ is the foil planform area and overline signifies cycle averaging.

A sample of the lift coefficient results are shown in Fig. 4 for both tail-like and flipper like kinematics across the range of sweep angles. The qualitative and quantitative features of the lift force over the cycle are relatively unaffected by doubling the sweep angle. In particular, our simulation results do not indicate a significant time delay in the peak lift with sweep, meaning that sweep is not changing the stability of the propulsive LEV. This conclusion is supported by the similarity in the LEV appearance on Λ20° and Λ40° in Fig. 3.

**Fig. 4.**
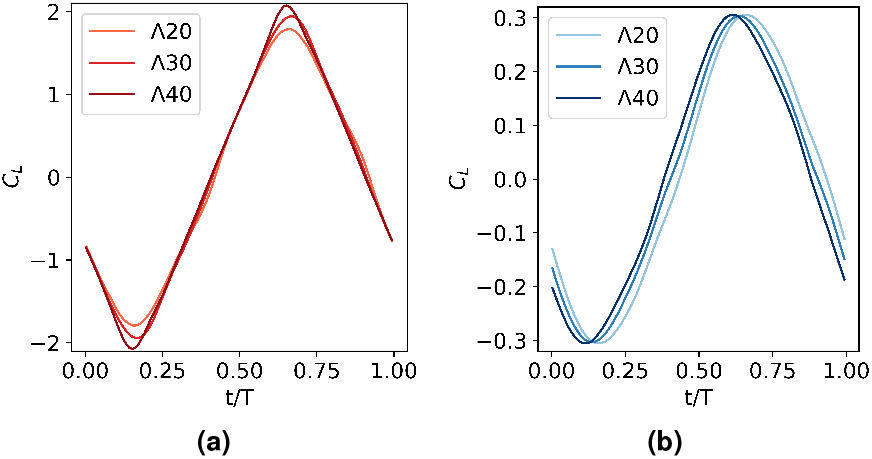
Lift coefficient of (a) tail-like and (b) flipper-like motion for 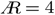, *St* = 0.3 and *k*^*\**^ = 0.3 in one cycle period (T). Both graphs show no LEV detachment delay as Λ increases.

The thrust coefficient and power also show a similar lack of influence of sweep on hydrodynamic performance. This information is presented in the supplementary material where we present the variation of mean thrust and lift coefficients (and their variance) as well as power coefficient (and its variance) and pitching moment for different 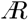 and kinematics. These simulation results suggest that sweep causes slight variations in the breakdown of the main vortex structures but this is not observed to have significant impact on cycle-to-cycle force and moment variations.

Fig. 5 summarises the influence of the geometric and motion parameters on the propulsive characteristics across 39 simulation cases and clearly illustrates our primary finding that sweep angle is not responsible for the large potential variation in flapping propulsion. Here, the geometric parameters are the sweep angle and 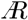, whereas the kinematic parameters are the motion type (heave, roll, etc) and flapping amplitude *A/l*^*^. Different *St* and *k*^*^ are achieved using two flapping amplitudes; a high amplitude, low frequency setting to mimic biological swimmer, and a low amplitude, high frequency setting to represent many robotic swimmers.

**Fig. 5.**
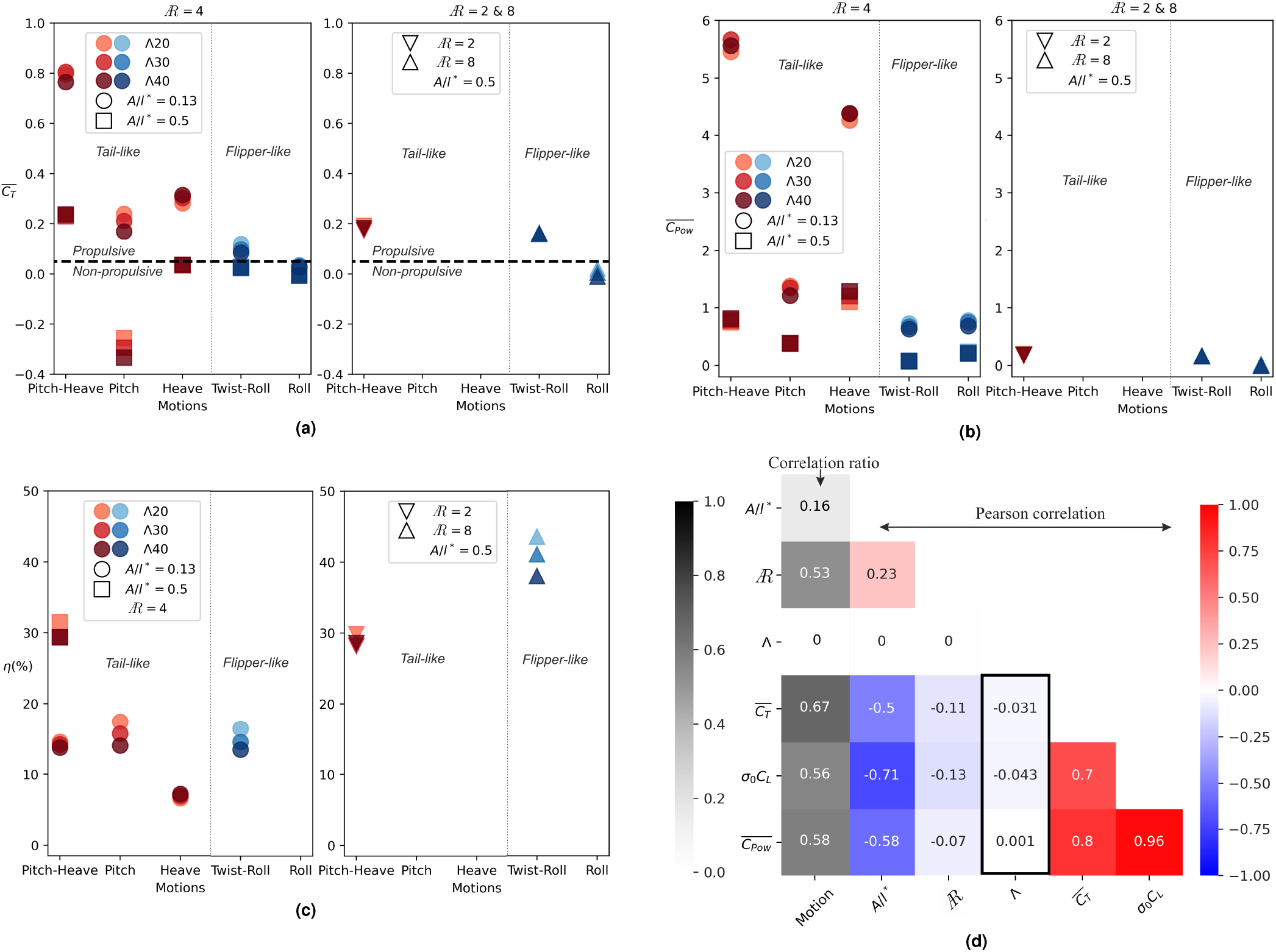
Results of 39 simulations are presented in different sweep angles Λ, aspect ratios 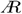, motions, and *A/l*^*\**^ for (a) mean thrust coefficients 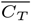, (b) mean power coefficients 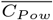 and (c) propulsive efficiencies *η* for the thrust producing cases 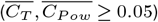. Correlation among the various input and output parameters for 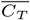, *η*, amplitude of the lift coefficient measured by the standard deviation *σ*_0_ *C*_L_, and 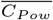 are presented in (d) a heat-map graph of correlation coefficients using Pearson and correlation ratio with a box highlighting Λ correlation. All graphs show that sweep angles have an insignificant impact on the output in spite of the large input variation convinced by the correlation values in the box of Fig (d). Please see an interactive graph in Supplementary Material for other input variations.

The cycle average thrust and power are shown in Fig. 5 (a,b). The motion type and amplitude produce large variations in these metrics (ranging from 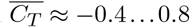 and 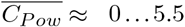) whereas the propulsive outputs are found to be substantially invariant to alterations of sweep angle and 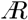. For positively propulsive cases 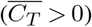, increasing sweep angle typically has an adverse effect on 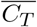; the largest being a 0.07 decrease between Λ20° to Λ40° for the engineering-inspired (low amplitude and intermediate 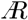) pure pitch cases. Compensating this loss of thrust, these cases also see a reduction of 0.17 in the required 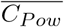 with increased sweep. In contrast, propulsive heave cases with engineering-inspired amplitude are seen to benefit from sweep angle with an increase of 0.03 in 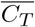, but with the corresponding increase of 0.12 in 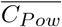. We also note that the propulsive tail-like motions produce relatively higher 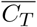 than the flipper-like motions and that biological-inspired amplitude cases generally have lower 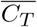 and 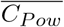 than engineering-inspired ones.

The propulsive efficiency is shown in Fig. 5 (c), which only includes configurations where 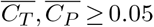 to avoid meaningless negative propulsive efficiencies and division by near-zero power (‘feathered’ conditions). As with the thrust and power, the impact of sweep angle on efficiency is found to be smaller than the motion parameters. The largest effects are a 5.6% efficiency decrease observed when increasing from Λ20° to Λ40° in the twist+roll (flipper-like) case for *A/l*^*^ = 0.5 and a 3.4% efficiency reduction for the pitch (tail-like) cases for *A/l*^*^ = 0.13. In contrast, we see that kinematics made up of a combination of degrees of freedom (twist+roll, heave+pitch) are much more beneficial than single degree-of-freedom motions (pure pitch, heave, or roll). The efficiencies above 25% belong to the cases with combined motions and higher amplitude (biological-inspired) conditions, with flipper-like motion giving the highest efficiency. This is caused by their lower 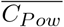 compared to engineering-inspired configurations. In addition, the penguin-like flipper case has a high *≈*40% efficiency which is achievable because the long span and large twist amplitude combine to achieve a similar maximum angle of attack along the span (30). All types of motion with lower amplitude (engineering-inspired) give efficiencies between 4%-20% with pure heave giving lower efficiencies because of higher 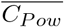.

Overall it is clear that kinematic parameters are more influential than the planform geometry for propulsive optimization. This can be summarized quantitatively using the correlations between the different input parameters and the propulsive outputs, Fig. 5(d). The Pearson’s correlation between the sweep angle and the propulsive statistics range from *−*0.043 to 0.001, whereas all the amplitude and motion parameters are strongly correlated with all the force and power coefficients, with levels of 0.50 to 0.71. The 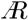 of the propulsive surface is also found to be secondary compared to the kinematics, with the largest absolute correlation of *−*0.13 between 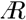 and the standard deviation of the lift coefficient.

## Conclusion

In conclusion, we have studied the influence of fin sweep angle and 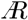 on flapping foil propulsive performance using a set of high resolution three dimensional viscous flow simulations. This enables us to more tightly control all other geometric and kinematic parameters than would be possible in biological or experimental studies, producing an extensive and focused data set of flow fields, integrated forces, and propulsive efficiencies. We have seen that sweep angle does not provide a significant advantage or disadvantage for tail-like propulsive foils. This can explain why biological data for sweep angle of fish caudal fins and mammal flukes have a large observed range and are not strongly correlated with kinematics or fin aspect ratio. From a biological perspective, this signifies that the so-called fastest fishes from Scombridae family or mammals from Delphinidae family in Fig. 1 can have either high or low sweep angle given any 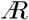 range without losing thrust and affecting their propulsive efficiency. Similarly, increased sweep angle does not give an advantage to flipper-like motions, instead tending to lose efficiency due to increased power requirements and loss of thrust. However, this loss is small, typically less than 10%, and can be compensated by increasing tip frequency, Strouhal number, and twist angle to recover the lost thrust force. The sweep angle also does not affect the lift coefficient, which is beneficial as flippers are also used as a lift regulator in animal swimming (31).

This study finds no benefit to sweep for steady-speed swimming, but does this does not preclude other more subtle potential fluid dynamic advantages. For example, in very high velocity motions more similar to animal maneuvering than steady swimming (32), experiments have observed a modulated *C*_*L*_ peak for Λ45° compared to Λ0° (21). In addition, previous simulations have show that fin-to-fin interactions may have an important effect on increasing propulsive efficiency (33). As such, is it still possible that sweptback angle may impart an advantage, despite not making a significant contribution to steady swimming efficiency. We do not address dynamic flexibility in this work, but we prove that the prescribed flexibility (twist) impacts greatly on the propulsive efficiency because it decreases side force *σ*_0_*C*_*L*_ and power coefficient 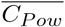. However, the sweep angle is still invariant to the twist, explaining why manta’s and cownose-ray’s flexible pectoral fins are found in the middle of sweep angle and 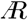 ranges in Fig.1 similar to most Scombridaes & Delphinidaes. This finding should help focus research in these more fruitful directions, and simplify the design of potential biologically-inspired vehicles and robots.

## Methods

The prescribed form of the heaving ℋ (*t*) and pitching *θ*(*t*) functions are given by

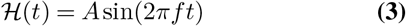

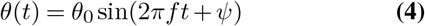

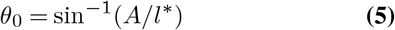

where *θ*_0_ the trailing-edge tip rotation amplitude, and the phase difference *ψ* = 90° are set based on the maximum performance for 2D motions (34). The twist is defined equivalently to pitch, with the amplitude increasing with distance from the root section. The pivot point of pitch and twist is at the leading edge of the root section. The roll motion is simply rotation about the inline axis, and the amplitude is set to achieve the same amplitude *A* at *l*^*^ similarly to maximum twist as described in (30).

The correlation data in Fig. 5 (d) use the Pearson coefficient of correlation between two continuous sets of data, and the correlation ratio between a category set and a continuous data set, defined as

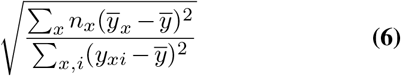

Where

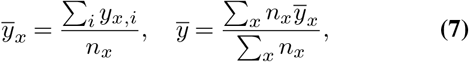

*x* category in each observation *y*_*xi*_, *i* index, *n*_*x*_ the number of observation in category *x*, and 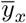 is the mean of category *x*. The flow simulation package used for this research is the Boundary Data Immersion Method (BDIM) (35). BDIM uses a robust and efficient Cartesian grid with an implicit LES (iLES) solver. BDIM uses analytic meta-equations for an immersed body in multi-phase flow with a smoothed interface domain using an integral kernel. BDIM has been validated, proved suitable for accurate force predictions on moving bodies such as flapping foils (36).

Symmetric conditions are enforced on both spanwise boundaries for the finite foil simulation. The domains extend from the pivot point to 4*C* at the front, 11*C* at the rear, 5.5*C* at the top and bottom. Meanwhile, the tip distance to the maximum spanwise domain is 3.2*C*. The grid convergence study provided in (19) using an infinite foil with a span length of 6*C* in *Re* = 5300 as a comparison of time-averaged thrust coefficient for different resolutions. The force coefficient converges to within 7% of the finer simulations (for 2D and 3D simulation) using a resolution of *C/*Δ*x* = 128. As a balance between the grid resolution and the number of simulations, this resolution is deemed sufficient to captures the dynamics of the flow.

## Supporting information

Statistics of tail-like and flipper-like motions for force coefficients, power coefficients, efficiencies, and pitching moments

The video corresponds to Figure 3 (a) and (c) in the manuscript. They present one-period phase evolution of instantaneous tail-like motion.

This video corresponds to Figure 3 (b) and (d) in the manuscript. They present one-period phase evolution of instantaneous flipper-like motion.

This is an interactive graph containing all data for Figure 5 in the manuscript.

## ACKNOWLEDGEMENTS

We would like to thank Indonesia Endowment Fund for Education (LPDP), IRIDIS High Performance Computing Facility with its associated support services at the University of Southampton and the Office of Naval Research Global award N62909-18-1-2091 for the completion of this work. We also want to thank Professor George Lauder (Harvard) for constructive discussions. Pertinent data will be made available upon publication.

