## Supplementary material for "Fin sweep angle does not determine flapping propulsive performance": Statistics of tail-like and flipper-like motions for force coefficients, power coefficients, efficiencies, and pitching moments

The tail-like kinematics exhibit insignificant changes in the statistics of the force-coefficients with the increase of sweep angle, (Fig. 1 (a)). There is a small increase in the range of lift coefficient (maximum and minimum $C_L$) as well as and the mean and maximum power. However, the thrust coefficients ($C_T$) show minimal change. This causes the efficiency ($\eta$), i.e. the ratio of average thrust to power, to decrease from 31% to 29% as the sweep angle increases. The flipper-like kinematics as presented in Fig. 1 (b) show the same tendency as the tail-like other than the mean $C_T$ which drops with increasing $\Lambda$. This condition makes the flipper-like motions lose $\approx 13\%$ efficiency between $\Lambda 20°$ and $\Lambda 40°$, while the tail-like motions only drop $\approx 2.2\%$.

We next check the combined significance of the sweep angle and different $\mathcal{AR}$. For tail-like motion, we halve the $\mathcal{AR}$ while maintaining the same $St$ and $k^*$ (Fig. 2). As a result, the statistic trends are even more insignificant than $\mathcal{AR} = 4$ especially the decrease in power coefficients, with an efficiency loss of only $\approx 1.7\%$. We use flipper kinematics to check the performance of long $\mathcal{AR} \approx 8$ foils since nearly flippers tend to have higher $\mathcal{AR}$ as shown in the manuscript Fig 1. In addition, turtles and penguins are observed to use higher frequency and higher twist amplitude (1, 2). This combination of kinematics is important because the observed large roll amplitude and frequency induces a high local angle of attack $\alpha_{max}$ towards the end of the foil, which would increase drag and reduce efficiency if applied alone. The increase in twist angle compensates for this, limiting $\alpha_{max}$ toward the foil tip to maintain high efficiency (3, 4). In our tests with $\mathcal{AR} = 8$ the large roll-angle amplitude $30° - 45°$ and high twist-angle amplitude $45° - 60°$ increase propulsive $\overline{C_T}$ while maintaining similar sectional maximum angle of attack $\alpha_{max}$. As such, these optimized flipper-like kinematics improve the efficiency, which now only drops $\approx 6\%$ for various sweep angle because $\overline{C_T}$ are almost constant.

We have checked the coefficients of moment for $\mathcal{AR}$ 2, 4 and 8, but there is also no significant alteration on the coefficients of pitching and rolling moment caused by the sweep-angle variation (see Tab. 1).

**Table 1. The effects of sweep angle ($\Lambda$) and aspect-ratio ($\mathcal{AR}$) variation on pitching moment coefficients( $C_{MZ}$) and their zero-mean standard deviation ($\sigma_0 C_{MZ}$) are negligible. (*) corresponds to those shown in the manuscript Figure 3 (a) and (c), and (**) to Figure 3 (b) and (d).**

| Motion | $\mathcal{AR}$ | $\Lambda$ (deg) | $C_{MZ}$ | $\sigma_0 C_{MZ}$ |
|---|---|---|---|---|
| *Pitch-Heave | 4 | 20 | 1.9e-4 | 5.2e-1 |
| | | 30 | -3.7e-4 | 5.6e-1 |
| | | 40 | -5.5e-5 | 5.6e-1 |
| Twist-Roll | 4 | 20 | 6.4e-5 | 9.6e-2 |
| | | 30 | -6.6e-6 | 1.0e-1 |
| | | 40 | -3.5e-5 | 1.1e-1 |
| Pitch-Heave | 2 | 20 | -1.3e-4 | 3.8e-1 |
| | | 30 | -2.1e-4 | 3.4e-1 |
| | | 40 | -1.5e-4 | 3.2e-1 |
| **Twist-Roll | 8 | 20 | -3.1e-4 | 2.4e-1 |
| | | 30 | 3.8e-4 | 2.7e-1 |
| | | 40 | -9.1e-4 | 2.7e-1 |

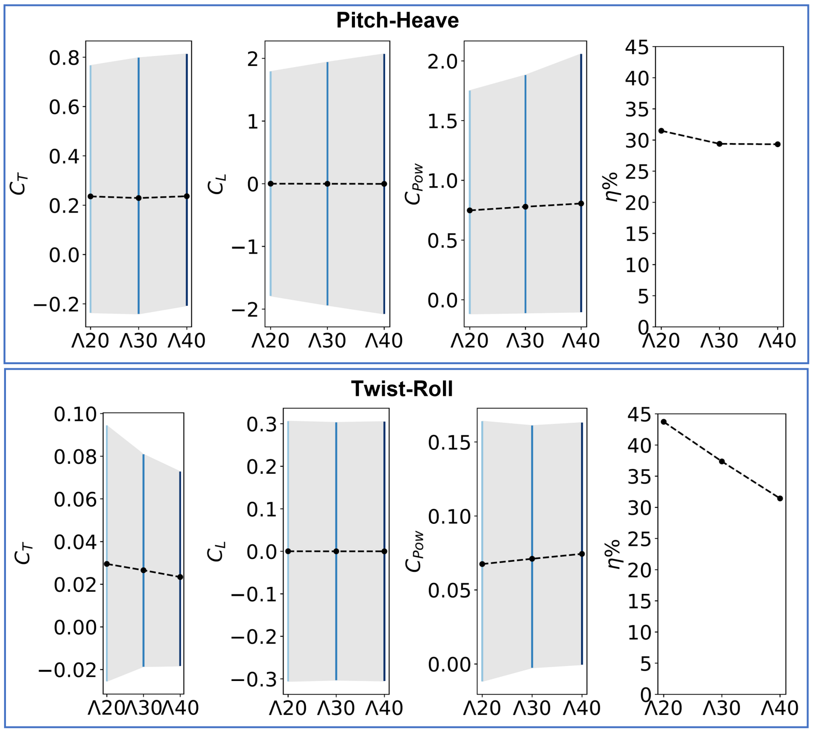


**(b)**

**Fig. 1.** Statistics of (a) tail-like and (b) flipper-like kinematics with the variation of sweep angle $\Lambda$ on $Æ = 4$ foil, $St = 0.3$, and $k^* = 0.3$. The tail-like cases in (a) correspond to those shown in the manuscript Figure 3 (a) and (c). Statistics are presented as the mean of coefficients of lift force $(C_L)$, thrust force $(C_t)$ and power $(C_{pow})$, shown in dashed lines, while grey shade represents the range the maximum and minimum values over a cycle. Force statistics are generally constant while a slight increase in power reduces the average efficiency $\eta$ when $\Lambda$ increases.

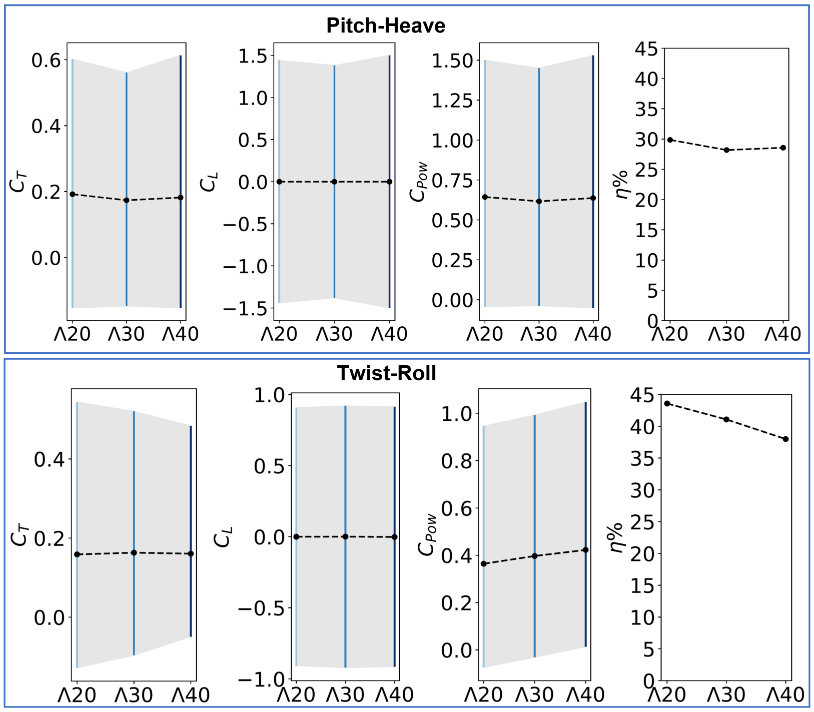


**(b)**

**Fig. 2.** Sweep angle variation even become more insignificant in different $Æ$ such as (a) lower $Æ = 2$ of tail-like in $St = 0.3$ and $k^* = 0.3$, and (b) higher $Æ = 8$, $St = 0.6$, and $k^* = 0.6$ of the flipper-like motion for optimum thrust and efficiency shown in the manuscript Figure 3 (b) and (d).
